# Iron promotes copper entry in *Streptococcus pneumoniae*

**DOI:** 10.1101/2023.12.17.572065

**Authors:** Yamil Sanchez-Rosario, Michael D.L. Johnson

## Abstract

Metals serve an important function at the host-pathogen interface, being used to leverage bacterial survival. To reduce bacterial viability in the host, some metals such as iron, are sequestered by the host, while others, such as copper are used to intoxicate bacteria. During infection, this serves the dual purpose of starving the bacteria of valuable resources while providing it with a toxic substance. By growing *Streptococcus pneumoniae,* a causative agent of multiple diseases including pneumonia, middle ear infections and sepsis, in the medium RPMI 1640 with a nanomolar concentration of iron, followed by exposure to a higher concentration of iron and copper, we observed an iron-dependent increase in copper association. This process was specific for iron and conserved in other *S. pneumoniae* serotypes. We performed single and double knockouts in selected iron transport systems and observed that under the same experimental conditions as wild-type strain, there was significantly less copper associated with the iron transport knockout bacteria. Taken together, we conclude that copper is inadvertently imported into the bacterial cell during iron acquisition.

## Introduction

The Gram-positive bacteria *Streptococcus pneumoniae* (pneumococcus) remains a formidable disease burden pathogen particularly in developing countries where it is responsible for 1.6 million cases of pneumonia and predominantly in children below the age of 5 [1]. Furthermore *S. pneumoniae* is also the causative agent of meningitis, middle ear and ophthalmic infections, endocarditis, and sepsis. One of this organism’s most prominent virulence factors is its neutral to negatively charged capsule. This amalgamation of sugar polymers that surrounds the bacteria is used to classify pneumococcus into serotypes, of which there are 100 serotypes described up to date. Capsule production is an energetically taxing endeavor requiring nutrient acquisition such as carbohydrates, polyamines, and metals.

Metals serve as protein cofactors and are crucial for proper bacterial function, growth, and multiplication. Existing in a host, exposes the bacteria to metal stress and challenges the bacteria to acquire necessary metals in a restrictive environment while avoiding the effects of metal intoxication. A central part of this sequestration and intoxication comes from the process of nutritional immunity, where the human host produces proteins that restrict the availability of nutrients such as metals. The interplay notably occurs with the bacteria’s need for iron juxtaposed with copper intoxication.

Nutritional immunity serves the dual purpose of protecting the host from dangerous redox damage and limiting the availability of metals for microbial use. Iron serves a crucial role in redox reactions and catalytic functions in mammalian and bacterial proteins. Some of the proteins that the human host uses to transport iron are hemoglobin and transferrin, while calprotectin and lactoferrin are used to restrict metals in microbial niches. It is estimated that the free available iron in the host is 10^-18^ M [2], making iron acquisition challenging for bacteria. Studies demonstrating the importance of iron for *S. pneumoniae’*s growth and survival show structural and growth rate deficiencies in iron limiting medium [3] [4], and conditional knockouts of iron import systems lead to virulence attenuation in murine models [5] [6]. Furthermore, exposure to host sources of iron, such as hemoglobin improves growth and changes in the pathogen’s transcriptome, causing differential expression of 145 genes two hours after exposure to hemoglobin, increasing the expression of virulence and colonization factors, and host-glycan degrading enzymes [7].

To obtain the necessary iron for survival some bacteria have developed passive and active strategies for iron acquisition. An active method of iron acquisition involves the production of cytotoxic proteins such as pneumolysin, which causes host-cell lysis and nutrient release from host cells. Additionally, during growth and metabolism, pneumococcus produces hydrogen peroxide [8]. Recently, this metabolic byproduct was shown to serve a biologically relevant function in iron acquisition via lysis and degradation of hemoglobin as a means for iron release [9], [10], [11]. During passive acquisition bacteria make small soluble compounds that diffuse into the environment and bind the necessary metals to deliver them to the bacterial cell. Siderophores are small iron scavengers made by some Gram-negative and Gram-positive bacteria inhabiting a close niche to the pneumococcus. *S. pneumoniae* is not reported to produce siderophores, but it has been known to utilize iron supplied by catecholate and hydroxamate siderophores in addition to iron from host carrier compounds such as transferrin, norepinephrine, hemin, and hemoglobin [12] [13] [14] [15], leading to increased bacterial growth and virulence expression. Pneumococcal iron scavenging is vital to survival in the host.

In *S. pneumoniae* TIGR4, the genes Sp_1032, Sp_1826 and Sp_1872 encode the substrate binding protein of three independent ABC-type importers responsible for iron acquisition (Figure 1, Table 1). All three importers have been reported to bind iron from siderophore-like molecules [4], [16]. In 2006 a survey of the distribution of Sp_1032 and Sp_1872 amongst pneumococcus and related commensal species found that these proteins were highly conserved between the pneumococcus [17]. The crystal structure of Sp_1032 shows a 1:1 molar ratio of either FeCl_3_ or Ferrichrome at a 2.1Å resolution [16]. Two glutamine residues from this substrate binding protein form salt bridges with arginines in the permease, thus allowing for iron movement into the cell.

**Figure 1.**
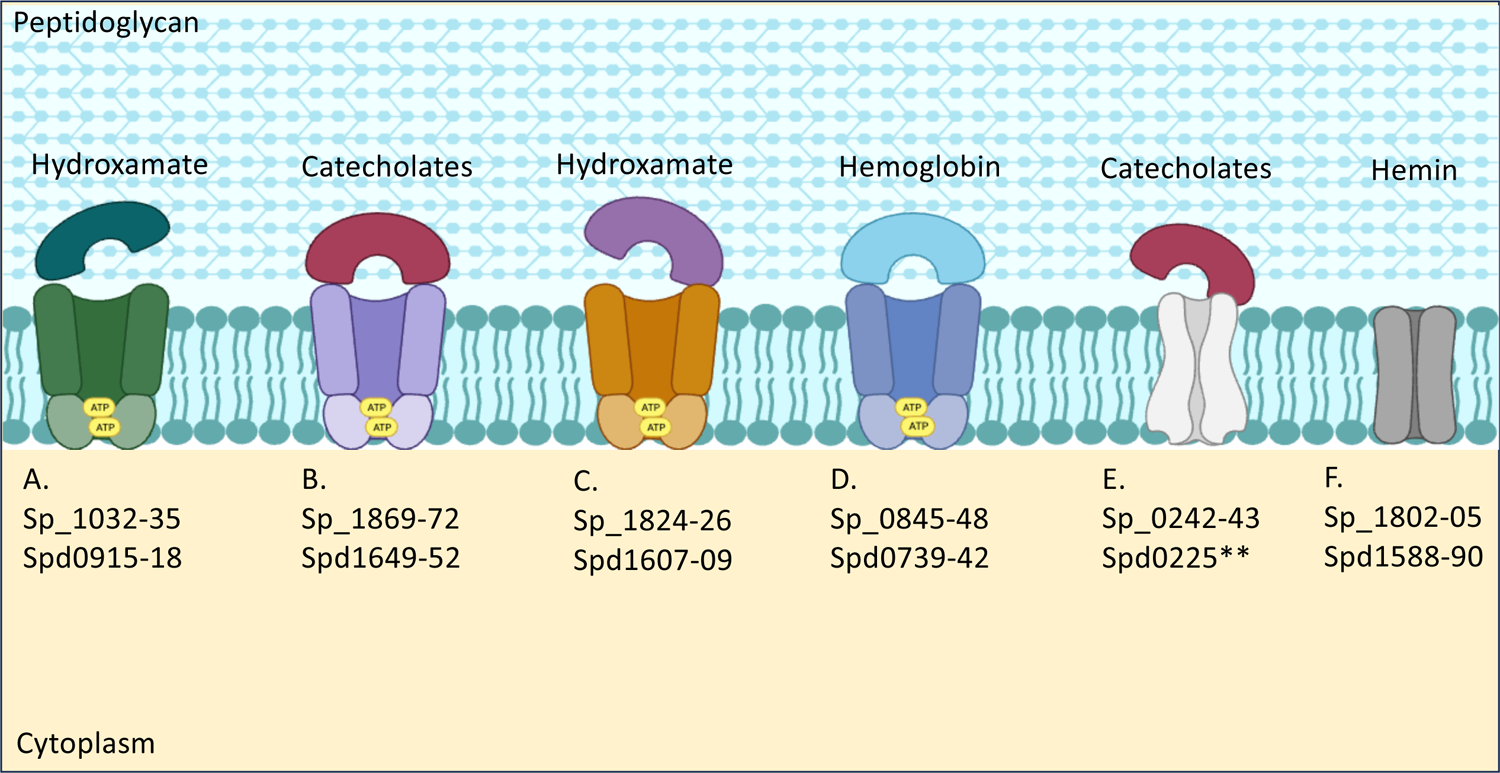
Described iron transport systems in *Streptococcus pneumoniae* TIGR4 and D39 gene loci annotation. Four systems belonging to the ATP Binding Cassette Family (A-C, E) and are reported to acquire iron facilitated through catecholate (enterobactin, norepinephrine) and hydroxamate (Ferrichrome) siderophores. One ABC system has been reported to bind hemoglobin (D). Two other systems are implicated in binding of catecholates or hemin (E-F) but appear to work in conjunction with a second transporter (sp_1869-70) to transfer iron to the cytoplasm. ** This annotation appears partly as a pseudogene in D39.

**Table 1.** Gene organization of ABC iron importers and protein size. ATP dependent iron transport in *S. pneumoniae* is arranged in clusters of 3-4 genes composed of a substrate binding protein, one or two proteins comprise the permease, and the ATPase.

| Gene locus | function | kDa |
| --- | --- | --- |
| Sp_1032 | SBP | 37.5 |
| Sp_1033-1034 | permease | 36-36 |
| Sp_1035 | ATPase | 29.6 |
| Sp_1869-1870 | permease | 35.9-36.2 |
| Sp_1871 | ATPase | 28.5 |
| Sp_1872 | SBP | 34.8 |
| Sp_1824 | permease | 62.6 |
| Sp_1825 | ATPase | 37.4 |
| Sp_1826 | SBP | 39.1 |
| Sp_0845 | SBP | 36.7 |
| Sp_0846 | ATPase | 55.1 |
| Sp_0847-48 | permease | 37.4-33.7 |

Structural dynamics of Sp_1872, using several methods including sequence analysis and NMR, agree with the conserved regions required for interaction with the substrate [18]. The authors further conclude that Sp_1872 is also able to coordinate bis and mono catechol complexes including the host norepinephrine. Normally, norepinephrine aid the host’s fight or flight response, but during infection, it acts as a siderophore, chelating iron away from transferrin or lactoferrin and becoming an accidental source of iron for bacterial [19], [18], [20].

There is no structural data on Sp_1826, however, one study that mutated other importers Sp_1872, Sp_1032 and Sp_0242 (Figure 1), found a substantial growth defect in this mutant. And the authors were only able to restore growth in iron depleted media after complementation with the Sp_1826 and the addition of Ferrichrome or FeCl_3_, suggesting the lipoprotein encoded by SP_1826 facilitated iron uptake from FeCl_3_ and Ferrichrome [6]. Ferrichrome is a xenosiderophore for *S. pneumoniae*, yet the bacteria can utilize iron supplied by this compound. To achieve this, pneumococcus employs the action of a substrate binding lipoprotein in association with a permease to deliver substrate to the intracellular space. Multiple reviews explaining these types of ABC transporters depict them with a c-style conformation or “venus fly” style of the protein, providing flexibility to allow binding of substrates [21], [22], [23], [24, 25]. Some of these lipoproteins are specific for the substrate bound and transported, while others can have certain promiscuity in binding and flexibility of transport which varies between systems and substrates. For example, the MtsA ABC transporter in *Streptococcus pyogenes* was first characterize as binding multiple substrates, such as Zn and Cu competing for the same site, and a second site with Fe (III) specificity [26]. In later works it was determined that the same system could also bind manganese [27].

Copper is another metal involved in nutritional immunity as it is necessary for growth, metabolism, and structural components of the mammalian host [28], however, it also has the potential to be toxic to the host. As such, homeostasis is achieved through the absorption of dietary copper, delivery through the body and excretion of excess copper. As a redox active metal, excess copper is toxic to bacteria via mismetallation, redox chemistry, and protein aggregation [29] [30]. As a response mechanism against the potential toxicity of copper, *S. pneumoniae* has evolved a copper export system that removes copper from the cell. Although we understand the mechanisms by which copper leaves the cell, the mechanism of entry remains elusive and is a major knowledge gap. The copper import system has not been identified or described and it is unlikely that there is a dedicated copper import system given the fact that no copper utilizing proteins have been identified in the pneumococcus. Therefore, the most likely explanation for the uptake of copper is through mismetallation of a different importer.

Traditional microbiological media contains sources of rich nutrients with polypeptides of undefined structure and concentrations, which studies show alters gene expression and can confound biological responses [3] [31, 32]. Furthermore, bacteria surviving at the host-pathogen interface will encounter host factors restricting its access to nutrient sources. Thus, to explore the complex network of metal interactions, including the mechanism of copper entry to the cell, we have chosen RPMI_mod_ 1640 [33] supplemented with nanomolar concentrations of trace metals as a defined media, which we considered more relevant to host conditions than traditional microbiological media. Under these more host-like restrictive conditions, we observed that during iron uptake, the pneumococcus inadvertently imported copper into the cytoplasm.

## Methods

### Bacterial growth

*S. pneumoniae* was routinely cultured overnight in tryptic soy agar plates supplemented with 5% sheep’s blood and incubated at 37°C with 5% CO_2_. Inoculums were taken from BAP or overnight cultures on RPMI_mod_ and inoculated onto RPMI_mod_ [33] supplemented with trace metals (246 µM MgCl_2_, 5.3 µM CaCl_2_, 179 nM FeSO_4_, 200 nM CuSO_4_, 173 nM ZnSO_4_, 87 nM MnCl_2_) and 0.1 mg/ml of catalase. Bacteria were grown to mid-exponential (OD600 nm 0.3-0.4) phase and the culture was divided into the different treatments (200 µM copper, 200 µM iron, 200 µM copper + iron) and control. Iron solutions were prepared fresh just before the addition step to avoid using oxidated iron for the treatments. Incubation of treatments was performed at 37°C and 5% CO_2_ for 30 minutes.

### Inductively Coupled Plasma Optical Emission Spectroscopy

Samples were placed in a ^-^3°C water bath to slow down metabolism, followed by two washes of cold buffer (Tris 50 mM, NaCl 150 mM, EDTA 200 mM at pH 7.6), and centrifugation 3500x g for 7 minutes at 4°C. Cold decanted samples were resuspended in 70% HNO_3,_ followed by overnight incubation at 65°C. After incubation, the samples were diluted to 2.5% HNO_3_ using milliQ water at 18.1 MΩ. As described above bacterial plate counts were performed in TSA + 5% Sheep’s Blood through serial dilutions.

Samples were analyzed for metal content using an iCAP PRO XDUO ICPOES with a wavelength 324.754 nm for copper, 213.856 nm for zinc, 259.373 nm for manganese and 259.940nm for iron. Standards were made using IV-STOCK-27 ICP calibration standard (Inorganic Ventures) and metal content of the washed samples was calculated using the Qtegra software.

### Bacterial quantification of living cells

Bacterial representative samples were taken from each of the treatments and serially diluted in PBS. Bacterial counts were performed in BAP to quantify the number of viable bacteria.

### Growth curves

Bacterial inoculums were seeded onto a 96-well plate to an optical density of 0.05. incubations at 37°C with 5% CO_2_. Measurements at 600nM were taken every 30 minutes after a 5-second orbital shake in a Cytation 5. Analysis was done using Prism 9 software from GraphPad.

### Pulse chase

The bacteria were grown to exponential phase and exposed for 30 minutes to either 200 µM copper, 200 µM copper + 200 µM iron or 200 µM iron. The cultures were then centrifuged and resuspended on fresh warm RPMI medium and incubated at 37°C for the appropriate time. Sampling for ICPOES was done at 30 and 90 minutes after media exchange.

### Gene knockouts

In-chromosomal deletion of the iron import genes Sp_1032-1035, Sp_1869-1872, Sp_1826-1824, and combinations were performed by antibiotic resistance replacement using either erythromycin or spectinomycin or a combination as the selective markers. Successful clones were sequence verified. The sequence reference used was from GenBank:AE005672.3.

### Statistics, figures, and graphics

Statistical analysis and figures were prepared using the software package GraphPad Prism 10. Graphics were created using Biorender.

## Results

The importance of metals in the growth and survival of bacteria is well established [34], [35], [36]. ABC transporters depend on the specificity of the substrate binding protein, but there is still promiscuity with these systems. Therefore, we explored the impact of biologically relevant divalent metal cations in the overall metal network of *S. pneumoniae*. We started by looking at metals requiring active transport and sought to understand how copper affected these interactions (Figure 2). After allowing bacteria to reach exponential growth (O.D.600nm 0.3-0.4), the culture was split and exposed to a combination of metals in conjunction with copper. We found that although zinc co-incubations increased the concentration of copper by two-fold, iron contributed far more to the increase in bacterial copper (Figure 2). We observed a 300-fold increase in copper over copper alone in bacteria exposed to a combination of copper and iron (Figure 2, Figure 3A). Because our media had a low concentration of metals and nutrients similar to the conditions encountered in the host environment, we postulated that the pneumococcus was iron starved. Thus, we supplied a higher concentration of iron during the growth period. Copper was still associated with the bacteria, albeit less than the more iron-depleted condition, but nevertheless, it suggested iron was an important modulator of copper entry (Figure 3B).

**Figure 2.**
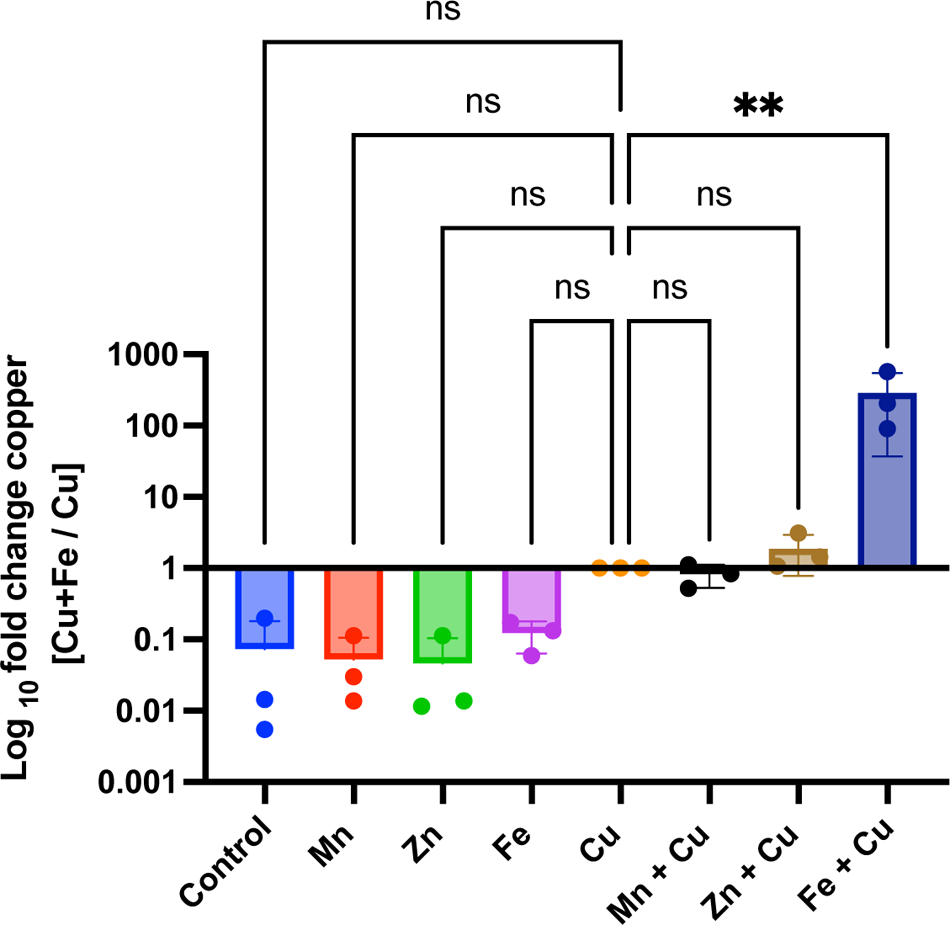
Other metals do not significantly increase the concentration of copper in TIGR4. Bacterial cells were grown on RPMI_mod_ with to a cell density of 1.3×10^8^ which correlates to 0.35-0.45 OD600. The culture was split into different treatments and exposed to 200 µM of indicated metal for 30 minutes. Samples were processed for ICPOES analysis. Copper concentration is indicated as fold change in metal combination over copper incubation. Report n=3 independent experiments. Statistical differences were measured in a one-way ANOVA (ns, non-significant; *, P < 0.05; **, P <0.01; ***, P <0.001; ****, P <0.0001).

**Figure 3.**
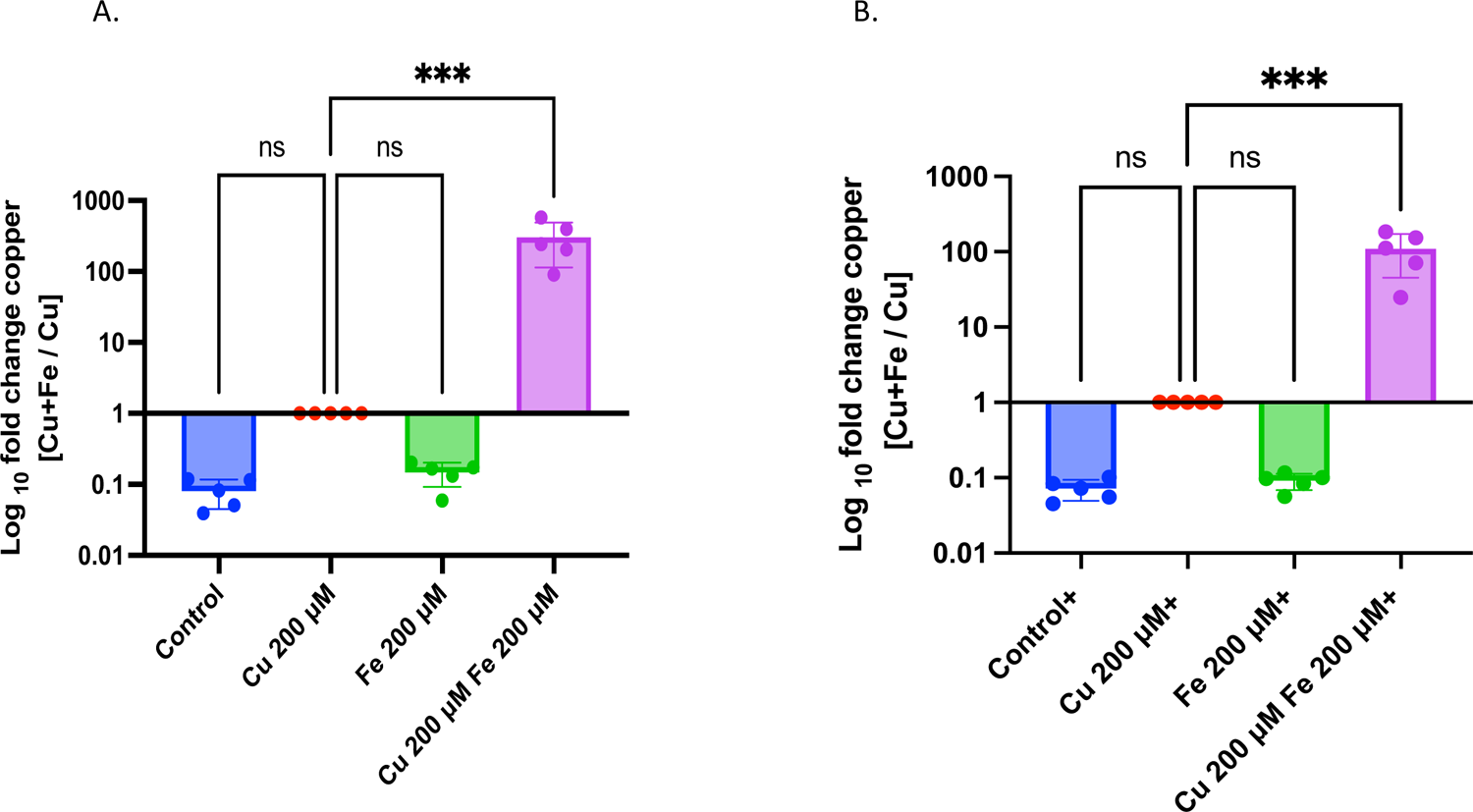
Iron drives copper association in TIGR4 cells. **A**. Bacterial cells were grown on RPMI_mod_ to a cell density of 1.3×10^8^ which correlates to 0.35-0.45 OD600. The culture was split into different treatments and exposed to 200 µM copper, iron, or combination for 30 minutes. **B**. Bacterial cells with added 100 µM FeSO_4_ were grown on RPMI_mod_ to a cell density of 1.3×10^8^ which correlates to 0.35-0.45 OD600. The culture was split into different treatments and exposed to 200 µM copper, iron or combination for 30 minutes. All samples were processed for ICPOES analysis. Copper concentration is indicated as fold change in metal combination over copper only incubation. Report n=5 independent experiments. Statistical differences were measured in a one-way ANOVA (ns, non-significant; *, P < 0.05; **, P <0.01; ***, P <0.001; ****, P <0.0001).

The pneumococcus is surrounded by a prominent antigen called a capsule, composed of an amalgamation of different sugar polymers. The combinations of sugars and other surface molecules create a relative negative charge of different strengths referred to as the zeta potential [37]. To determine if the capsule charge contributed to the copper signal observed during ICPOES, we selected different capsule compositions based on their zeta potential and a non-encapsulated isogenic mutant [38]. Based on the Chun *et al*. 2023 work, the relative zeta potential of chosen strains were as follows: 33F most neutral with −5 mV, followed by the TR, an unencapsulated mutant with −11 mV, D39 a type 2 capsule with −12 mV, TIGR4 a type 4 capsule with −16 mV and the most electronegative 16F with −22 mV. We found that under the same conditions, copper was still associated with the bacteria in the presence of iron independent of the capsule serotype or zeta potential (Figure 4). Furthermore, the capsule mutant also had significantly more copper in the presence of iron than copper alone, suggesting that the increase in copper is not associated with the capsule itself (Figure 4). Taken together, copper is not associated with the outside of the bacteria as the increase occurs only in the presence of iron and is capsule-independent.

**Figure 4.**
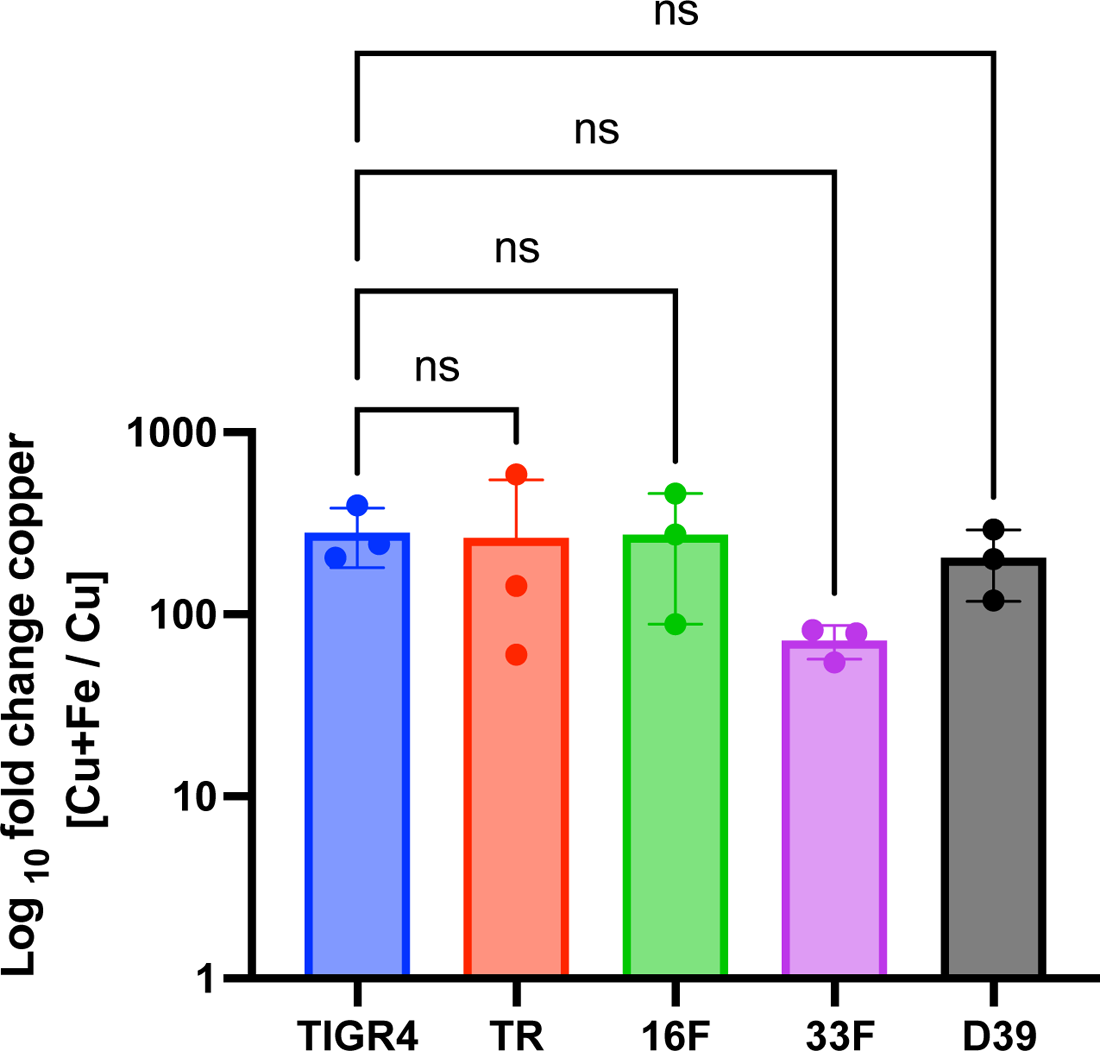
Capsule does not contribute to copper increase. Copper also increases in Fe 200 µM + Cu 200 µM combination treatments in other capsule serotypes suggesting capsule does not mediate copper association. Report n=3 independent experiments. Statistical differences were measured in a one-way ANOVA (ns, non-significant; *, P < 0.05; **, P <0.01; ***, P <0.001; ****, P <0.0001).

While the copper export system leads to a decrease of copper associated with bacteria over time, the speed and specificity with which this increased level of internal copper is exported is unknown. We performed a pulse chase experiment to determine if copper ultimately leaves the cells. While increased copper uptake coincided with the presence of iron, we tested if the association was due to a lack of copper export. By removing metal exposure and performing a media exchange, the levels of copper begin to decrease, and 90 minutes after media exchange, the levels of copper have dropped significantly in the metal combination treatment, indicating copper is exported from the cell (Figure 5). As a control, we also looked at the concentration of iron in the pellet during the same time points as copper. Iron remains constant after media exchange, and resuspension on low iron media does not change the concentrations associated with the cells (Supplemental Figure 1). Taking these data together, the decrease in copper and no change in iron after media exchange suggests the decrease in copper concentrations is associated with copper export and not with loss of pellet through lysis.

**Figure 5.**
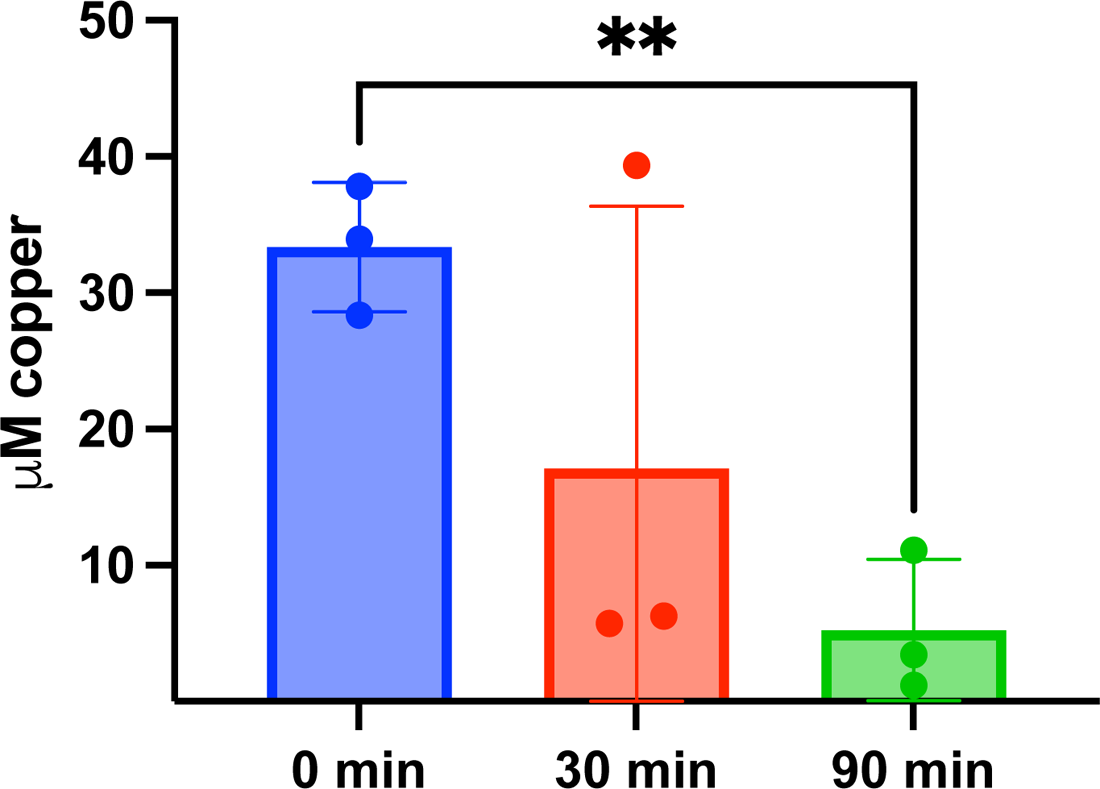
Copper concentration trends downward after removal of metal stress. TIGR4 cells were grown on RPMI_mod_ and exposed to single metals or in combination of 200 µM copper + 200 µM iron for 30 minutes. Bacteria was centrifuged and resuspended on fresh prewarmed RPMI_mod_, followed by incubation. Samples were collected at 30 minutes or 90 minutes after resuspension. Report n=3 independent experiments. Statistics using an unpaired T-test (ns, non-significant; *, P < 0.05; **, P <0.01; ***, P <0.001; ****, P <0.0001).

Having identified that iron import is a potential source for copper entry into the cell, we then decided to knockout some of the iron import systems (Figure 1). We chose to knock out single import systems (Sp_1032-35, Sp_1869-72 and Sp_1824-26) as well as double combinations (Sp_1032-35 + Sp_1869-72 or Sp_1824-26 + Sp_1869-72) to explore their individual systematic contributions. We focused on the primary ABC importers as their ability to bind iron from siderophores suggested an innate flexibility that could result in a source of mismetallation. We decided against triple knockouts as previous literature shows these mutants have a growth defect [4]. Given that our growth media is more restrictive in nutrients when compared to traditional microbiological growth media, and metals are kept at a nanomolar concentration compounded to the available iron form, it was expected and observed that the iron import mutants have a longer lag phase when compared to the parental strain (Supplemental figure 2).

However, because we collected our samples during exponential phase, at which point cells are actively growing, we don’t expect our results to be influenced by the longer lag phase. Figure 6 A shows that iron import mutants have a significant decrease in intracellular iron per CFU after 30 minutes of 200 µM FeSO_4_ exposure compared to the parental strain. Likewise, these mutants also have significantly less copper associated per CFU after copper only exposure, suggesting the mechanism of entry has now been restricted by reducing the number of potential entry points to the cell (Figure 6 B).

**Figure 6.**
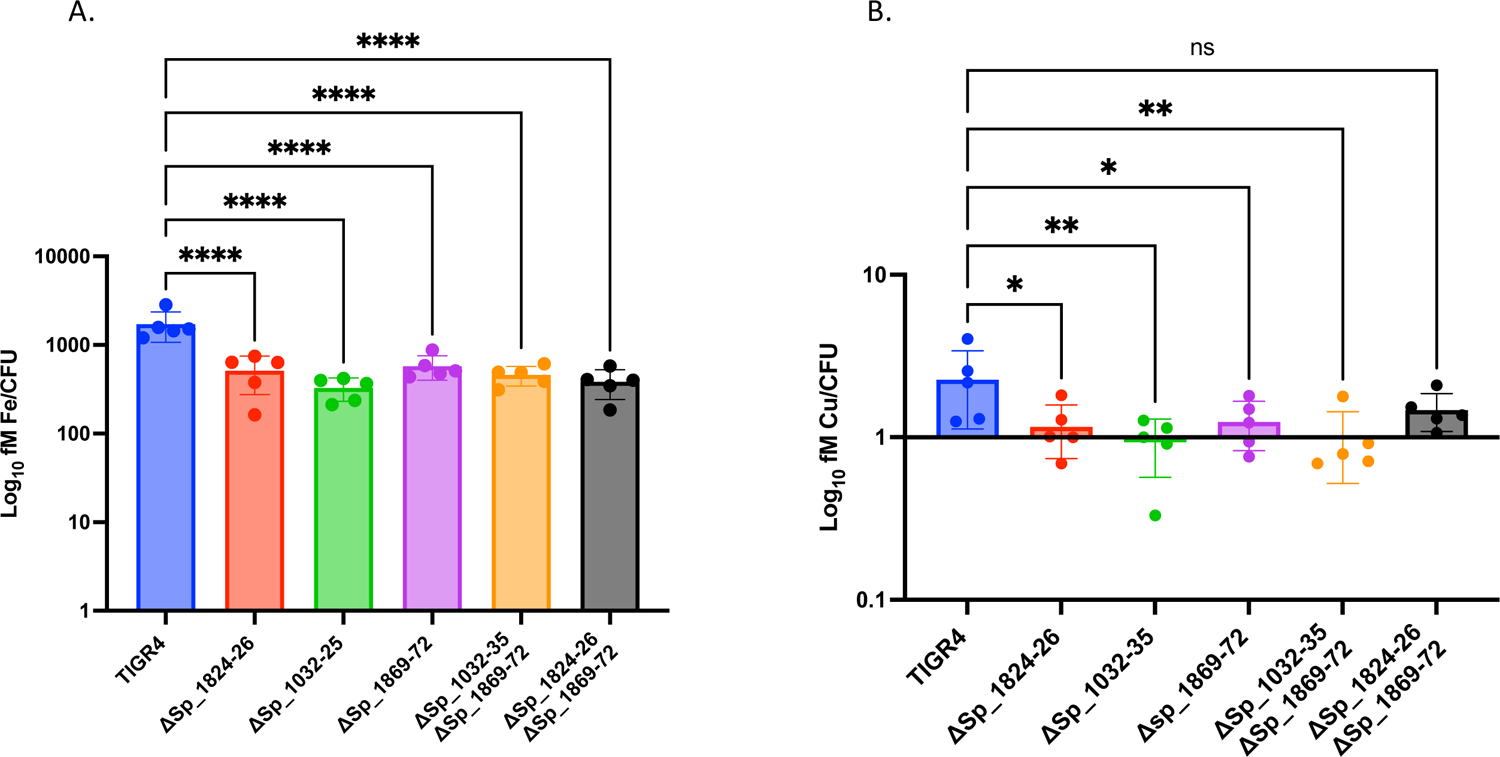
**A**. Iron concentration decreases in iron transport knockouts as compared to the parental strain. Bacterial cells were grown on RPMI_mod_. The culture was split into different treatments and exposed to 200 µM copper, iron, or combination for 30 minutes. Samples were processed for ICPOES analysis. Iron is reported as fM/CFU in the iron only sample. **B. Copper concentration decreases in iron transport knockouts as compared to the parental strain**. Bacterial cells were treated as in A. Copper concentration is reported as fM/CFU in the copper only sample. Report n=5 independent experiments. Statistical differences were measured in a one-way ANOVA (ns, non-significant; *, P < 0.05; **, P <0.01; ***, P <0.001; ****, P <0.0001).

Furthermore, in copper + iron combination treatment, we found that single iron import knockouts have copper concentrations that are significantly lower when compared to the parental strain under the same conditions, further implicating that iron importers are involved in the uptake of copper (Figure 7).

**Figure 7.**
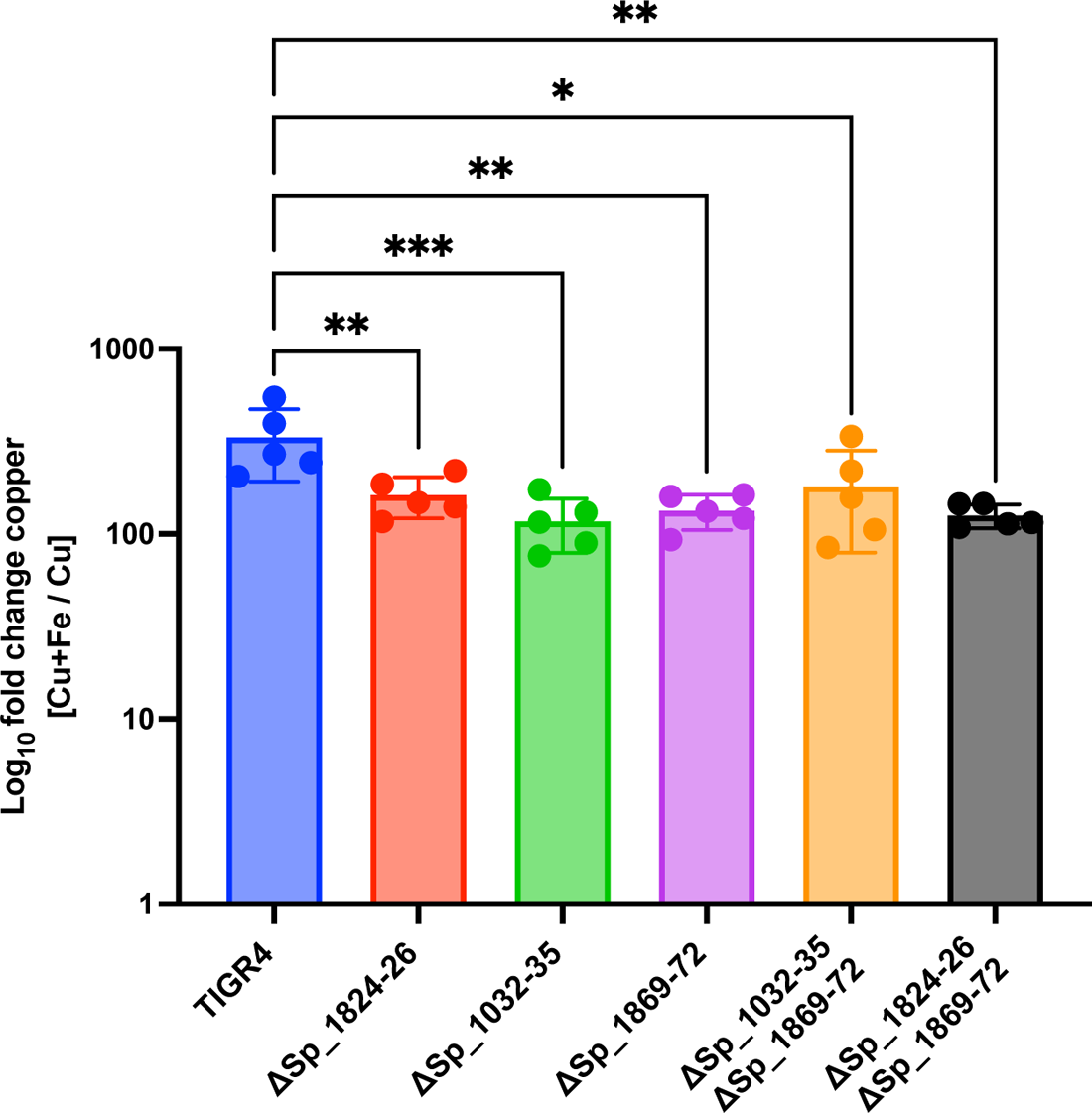
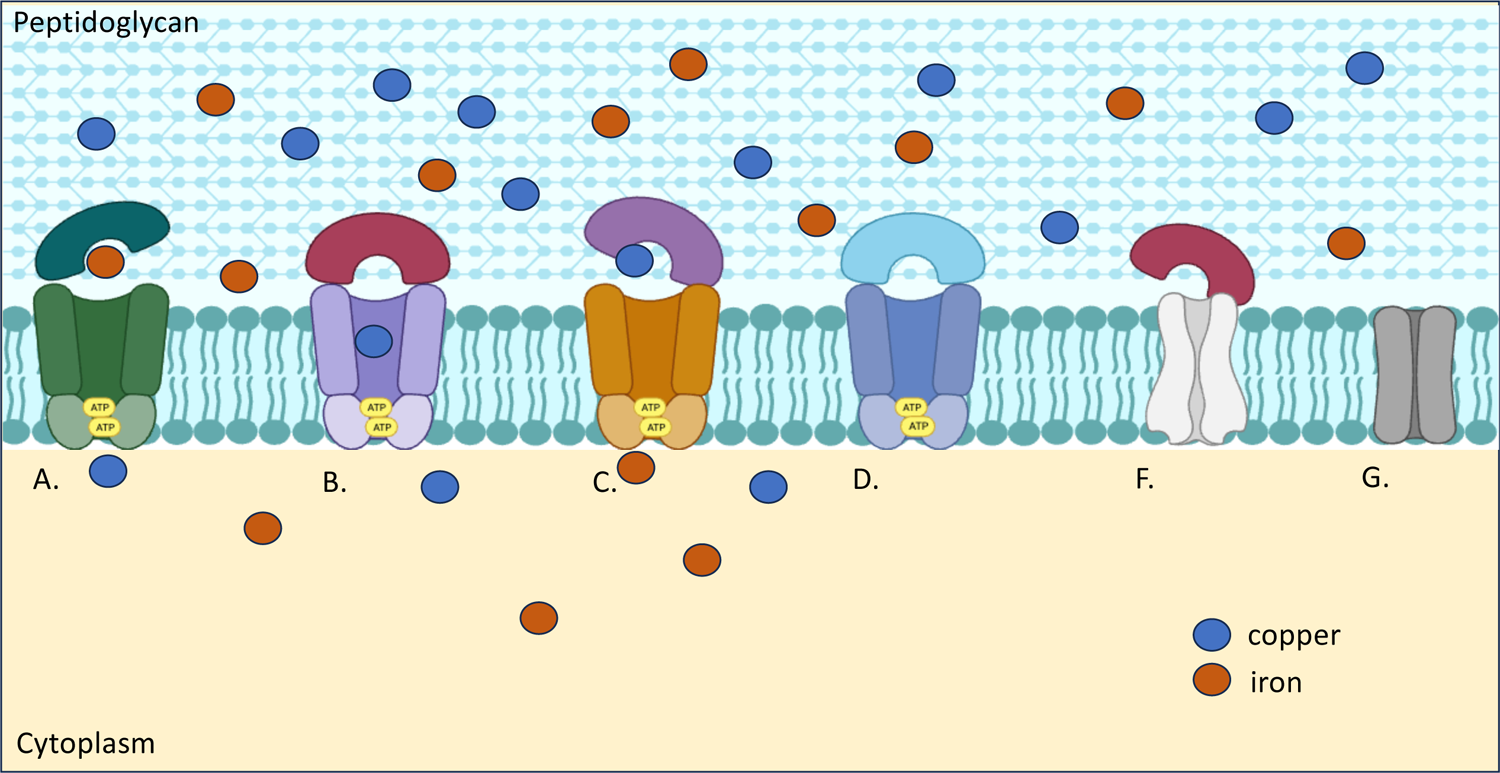
Copper concentration decreases in iron transport knockouts as compared to the parental strain. Bacterial cells were grown on RPMI_mod_. The culture was split into different treatments and exposed to 200 µM copper, iron, or combination for 30 minutes. Samples were processed for ICPOES analysis. Copper concentration is indicated as fold change in combination over copper incubation. Report n=5 independent experiments. Statistical differences were measured in a one-way ANOVA (ns, non-significant; *, P < 0.05; **, P <0.01; ***, P <0.001; ****, P <0.0001).

## Discussion

Traditional microbiological growth media can significantly impact gene expression and mask important phenotypes, such as artificially increasing the resistance to environmental stresses in vitro. To prevent confounding biological effects brought about by nutrient-dense media, we chose modified RPMI 1640 supplemented with nanomolar concentrations of trace metals to investigate the interconnection of metals in the biology of *S. pneumoniae*. Under these conditions, we observed an increase in intracellular copper concentration that was dependent on the presence of iron. Since iron was kept at nanomolar concentration in the growth medium, we hypothesized that the pneumococcal iron uptake systems that satisfy iron requirements during growth also uptake copper via mismetallation of these import systems.

To determine if other metals could promote the association of copper with the bacteria, we also performed coincubations with Zn or Mn. We chose these metals as opposed to other metals as they are known to have active transport, biological relevance, and low bioavailability in the host. We chose to keep the concentration of metal consistent between different metal treatments and acknowledge that except for zinc, the other metals tested were outside (higher) the biological range for either homeostatic or infection conditions [39], [40]. Regardless, neither zinc nor manganese provided at similar levels as iron seemed to significantly affect the levels of intracellular copper. This observation may be attributed by the pneumococcus having fewer zinc and manganese importers, as compared to iron importers and, manganese and zinc exporters where there are no iron exporters in *S. pneumoniae*.

We also explored the possibility that the increase in copper was explained by electrostatic interactions of the divalent metal cations with the negatively charged bacterial capsule or by nonfunctional copper export. We found that there was copper association even in the absence of capsule and that copper export still occurred, thus implicating iron importers as the culprits for iron import. This hypothesis was supported by observing significant reductions in copper, both independent and given concurrently with iron in the iron importer deletion mutants.

In the Irving Williams series, copper holds a more stable complex with enzymes when compared to other first row transition metals. Therefore, if we apply this principle to the transport of metals, copper could mismetallate substrate binding proteins of metal importers. Evidence supporting mismetallation of substrate binding proteins exists. The pneumococcal manganese binding protein PsaA also binds zinc with a similar type of coordination chemistry but different kinetics [41]. However, *in vitro* experiments indicate that the protein cannot release Zn upon EDTA treatment, suggesting the incorrect metal keeps the protein in the lock position preventing transport of the wrong metal [42]. Interestingly, in *S. pyogenes*, copper has previously been shown to downregulate both the zinc and manganese systems [43].

Another example is the Gram-negative *Escherichia coli*, in which the protein NikA binds nickel, cobalt and copper with similar coordination chemistry. However, thermodynamic studies show that only nickel transport is favorable [44]. Work using differential scanning fluorimetry looking AdcAII, the zinc transporter in *S. pneumoniae* demonstrated Zn, Co, Mn, Ni and Cu were able to induce significant changes in protein melting curves [45], suggesting that transition metals have the ability to interact with this protein, albeit to what extent it leads to functional transport remains unknown. From these *in vitro* data, mismetallation of ABC transporters leads to non-functional systems and rejection of the non-cognate metal. However, *in vitro* experiments are unlikely to account for the different redox states faced by the proteins as they transport the metals from an oxidized environment to a more reduced environment and the oxidation state of the metal can affect the recognition by a protein [46]. Furthermore, physiological relevant conditions such as the presence of bicarbonate later demonstrated that the lipoprotein MtsA of *S. pyogenes* has distinct affinities from what was originally reported from *in vitro* experiments [26]. The presence of bicarbonate influences the hierarchy or divalent metal cations binding to MtsA now showing higher affinity for ferrous iron, Fe^2+^> Fe^3+^> Cu^2+^> Mn^2+^> Zn^2+^ [47], indicative of how other factors can influence oxidation state and or metal predilection during metal acquisition.

Recently, a mismetallation of an ABC transporter was demonstrated in *Staphylococcus aureus* [48]. This bacterium colonizes the nasal cavity in the human host and is subjected to metal restriction. *S. aureus* is a Gram-positive bacterium which produces metal chelators for zinc acquisition under the low zinc environment induced by the host. This metal chelator called Staphylopine works with a metallopore system CntE to uptake and transport zinc into the cell. However, it was found that under zinc depleted conditions and in the presence of excess copper, Staphylopine non-specifically binds and transport copper into the cell leading to copper intoxication. Another example of transporter mismetallation was recently reported by Akbari *et al*. while investigating the pathogenesis of Group B Streptococci in diabetic wound infections. In their short communication they report the possibility of a nickel importer promiscuously introducing copper into the cell as their nikA mutant possessed significantly lower concentrations of copper [49].

In summary, our evidence suggests copper enters the pneumococcus via iron import systems likely via mismetallation. Since there are six known iron transport systems in the pneumococcus but fewer systems for transporting other metals, it is reasonable to interpret the larger increase in copper to promiscuous entry of copper through the multiple iron importers in the cell rather than a smaller contribution from other metal transporters such as Mn or Zn. Future directions will explore the mechanism of copper transport using biophysical techniques to identify copper coordination through the iron transporters.

---

**Supplemental Figure 1.** Iron remains constant in TIGR4 cells pulse chase experiments. TIGR4 cells were grown on RPMI_mod_ and exposed to single metals or in combination of 200 µM copper + 200 µM iron for 30 minutes. Bacteria was centrifuged and resuspended on fresh prewarmed RPMI_mod_, followed by incubation. Samples were collected at 30 minutes or 90 minutes after resuspension. Report n=3 independent experiments. Statistics using One-Way ANOVA (ns, non-significant; *, P < 0.05; **, P <0.01; ***, P <0.001; ****, P <0.0001).

**Supplemental Figure 2.** Growth curve comparison of iron import mutans and TIGR4 parental strain on RPMI_mod_. Mutants display a longer lag phase in comparison to the parental strain. Model of copper entry through mismetallation of the iron import systems. Multiple iron import systems are recruited to satisfy the necessary iron required by *S. pneumoniae* to grow and multiply in the low iron environment. Once the transporters are exposed to excess copper in the presence of iron, the importers (A-C were tested, is unknown for D-G) will uptake both iron and copper. At this moment the kinetics and or stoichiometry by which this process occurs is unknown.

