## Supplementary figures and images for "Iron promotes copper entry in *Streptococcus pneumoniae*"

### Supplemental figures

Supplemental Figure 1.

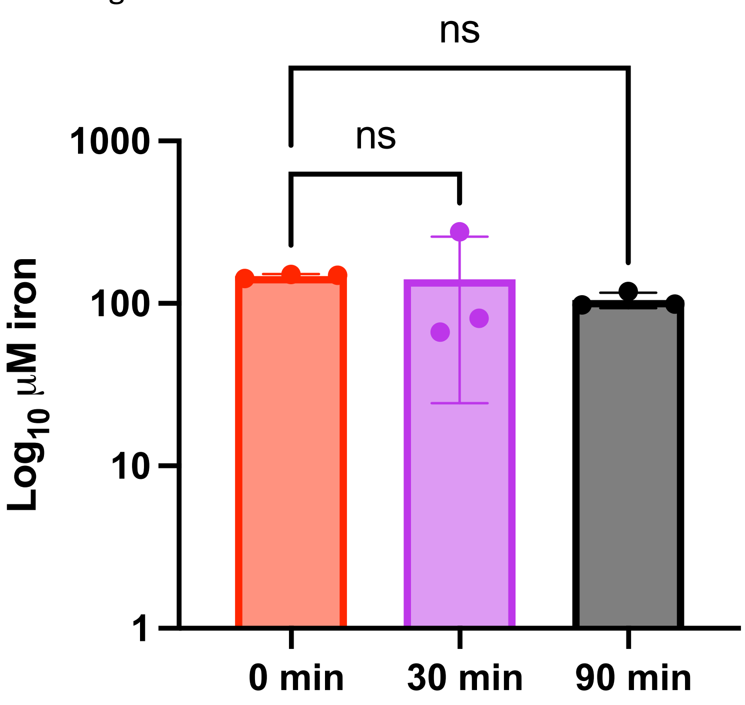

Supplemental Figure 2.

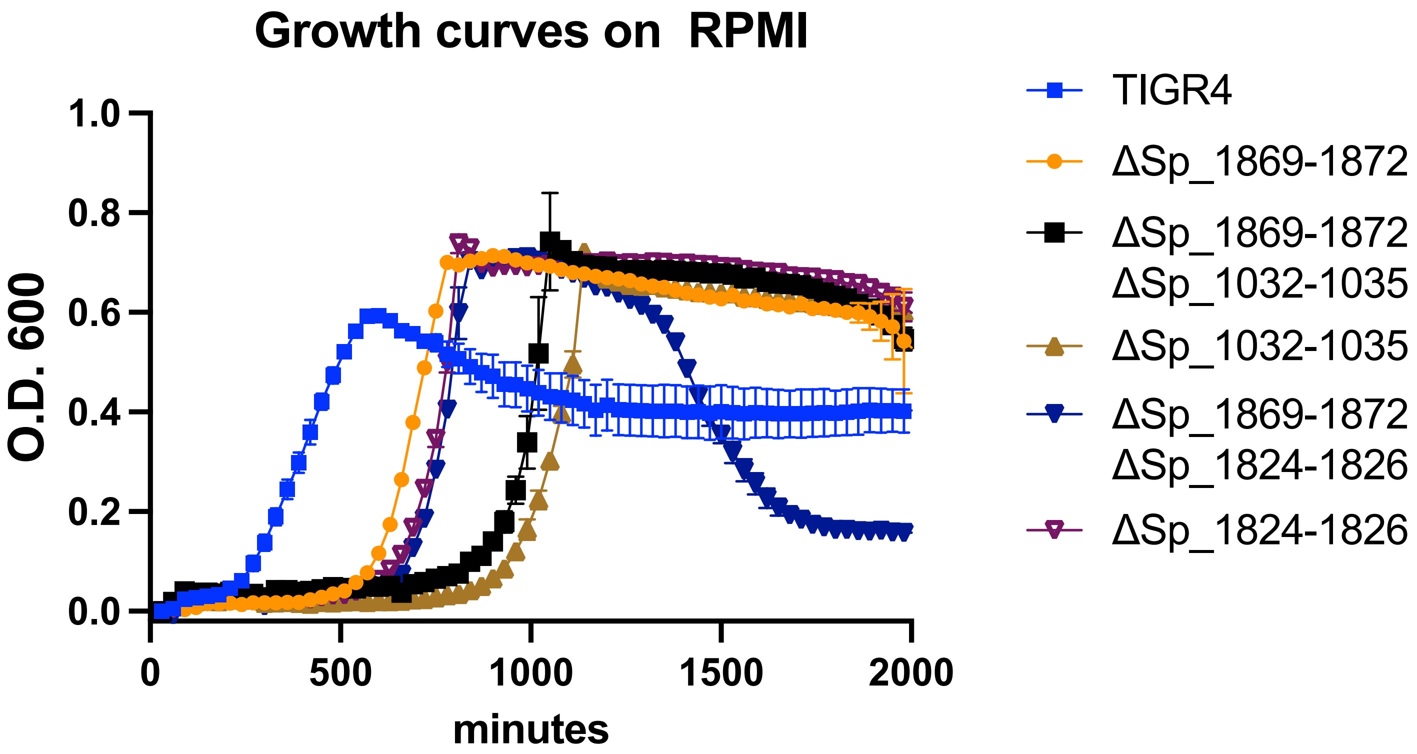
